# mCLAS adaptively rescues disease-specific sleep and wake phenotypes in neurodegeneration

**DOI:** 10.1101/2024.08.20.608757

**Authors:** Inês Dias, Christian R. Baumann, Daniela Noain

## Abstract

Sleep alterations are hallmarks of prodromal Alzheimer’s (AD) and Parkinson’s disease (PD), with fundamental neuropathological processes of both diseases showing susceptibility of change upon deep sleep modulation. However, promising pharmacological deep sleep enhancement results are hindered by specificity and scalability issues, thus advocating for noninvasive slow-wave activity (SWA) boosting methods to investigate the links between deep sleep and neurodegeneration. Accordingly, we have recently introduced mouse closed-loop auditory stimulation (mCLAS), which is able to successfully boost SWA during deep sleep in neurodegeneration models. Here, we aim at further exploring mCLAS’ acute effect onto disease-specific sleep and wake alterations in AD (Tg2576) and PD (M83) mice. We found that mCLAS adaptively rescues pathological sleep and wake traits depending on the disease-specific impairments observed at baseline in each model. Notably, in AD mice mCLAS significantly increases NREM long/short bout ratio, decreases vigilance state distances by decreasing transition velocities and increases the percentage of cumulative time spent in NREM sleep in the last three hours of the dark period. Contrastingly, in PD mice mCLAS significantly decreases NREM sleep consolidation, by potentiating faster and more frequent transitions between vigilance states, decreases average EMG muscle tone during REM sleep and increases alpha power in WAKE and NREM sleep. Overall, our results indicate that mCLAS selectively prompts an acute alleviation of neurodegeneration-associated sleep and wake phenotypes, by either potentiating sleep consolidation and vigilance state stability in AD or by rescuing bradysomnia and decreasing cortical hyperexcitability in PD. Further experiments assessing the electrophysiological, neuropathological and behavioural long-term effects of mCLAS in neurodegeneration may majorly impact the clinical establishment of sleep-based therapies.

## Introduction

Sleep-wake disturbances are a common feature of the asymptomatic preclinical stages of Alzheimer’s (AD) and Parkinson’s disease (PD), suggesting sleep may play a crucial role in the onset and progression of neurodegenerative processes (Holth et al., 2019; J. E. Kang et al., 2009; Lucey, 2020; Minakawa et al., 2017; Owen & Veasey, 2020; Qiu et al., 2016).

Hallmarks of early-stage AD include altered vigilance state patterns, as decreased time spent in NREM sleep, increased NREM sleep fragmentation and insomnia (Bubu et al., 2017; Lim et al., 2013; Sindi et al., 2018). Furthermore, increased locomotor activity prior to sleep onset together with increased agitation, aggression, confusion and emotional behavioural disruptions during early evening hours, are collectively described as the sundowning circadian syndrome in AD patients (Todd, 2020).

Contrastingly, PD features a generalized slowing of patients’ EEG signal and abnormalities in spectral EEG frequencies (termed bradysomnia), which correlate with anxiety, cognitive decline and apathy (Cozac et al., 2016; Imbach et al., 2016; Soikkeli et al., 1991; Zhu et al., 2019). Moreover, REM sleep behaviour disorder (RBD), characterized by the loss of normal skeletal muscle atonia during REM sleep, is also described as an important underpinning of early-stage PD (Berg et al., 2021; Postuma, 2014; Schenck et al., 1986).

Consequently, a bidirectional relationship between sleep and neurodegeneration is acknowledged, with neurodegeneration progression worsened by deep sleep disruption (Ju et al., 2014; Mapelli, 2015; Schreiner et al., 2019; Wang & Holtzman, 2019). Importantly, pharmacological sleep induction protocols even though effective at increasing deep sleep in animal models of AD and PD (J. Kang et al., 2009; Morawska et al., 2021) and in patients (Büchele et al., 2018; Mednick et al., 2013; Walsh, 2009), present major practical limitations as high risk of tolerance, dependency, misuse and lack specificity (Cellini & Capuozzo, 2018; Lie et al., 2015). Therefore, there is a clear need to develop and employ nonpharmacological and noninvasive tools to specifically enhance deep sleep in neurodegeneration (Wilckens et al., 2018).

In this context, closed-loop auditory stimulation (CLAS) appears as a promising approach to modulate sleep slow-waves in a precise manner, enhancing deep sleep when auditory stimuli are targeted to the up-phase of slow-waves, with improved memory recall and amyloid dynamics in older adults and patients with mild cognitive impairment (Papalambros et al., 2017, 2019; Wunderlin et al., 2023, 2024; Zeller et al., 2024). This technique has been regarded as safe, reliable and noninvasive to apply in a home-based environment, which may be ideal for patients displaying cognitive or motor deficits (Ferster et al., 2019; Lustenberger et al., 2022). Despite these remarkable prospects, the lack of animal models allowing the combination of phase-targeted CLAS in a neurodegeneration disease context has impaired its optimization and translation into clinical settings.

To bridge this gap, we have recently established mouse CLAS (mCLAS) in two mouse models of neurodegeneration, showing that phase-targeted mCLAS can be successfully implemented in healthy and diseased mice in a precise manner (Dias et al., n.d.). Moreover, preliminary data showed mCLAS acutely alleviate macrostructural sleep disturbances such as increased NREM sleep fragmentation, but further in-depth assessments are needed to infer whether this translational technique may be beneficial in rescuing disease-specific sleep alterations in early stages of AD and PD neurodegeneration.

Here, we assessed mCLAS’ short-term effects onto disease-specific sleep and wake disturbances in two mouse models of neurodegeneration: 7-month old AD mice, Tg2576 line (Hsiao et al., 1996); and 7-month old PD mice, M83 line (Giasson et al., 2002). Our aims comprised assessing the stability and dynamics of vigilance states (WAKE, NREM sleep and REM sleep) and their transitions in both models and probing mCLAS’ effect onto vigilance state patterns in AD mice, as well as altered muscle atonia, spectral power, general EEG slowing and hyperexcitability in PD mice.

## Methods

### Animal models and surgeries

We used the Tg2576 model of AD (Hsiao et al., 1996), overexpressing a mutant form of the human amyloid precursor protein (APP) gene, the APPK670/671L; and the M83 model of PD (Giasson et al., 2002), expressing the A53T-mutated variant of the human α-synuclein gene under the direction of the mouse prion protein promoter. The study design included female and male transgenic (TG) mice of each model (AD: n = 7, 4 females; and PD: n = 7, 6 females) at a prodromal disease stage (6.5 months-of-age at implantation, plaque- or heavy inclusion-free) and their wild-type (WT) littermates (WT_AD_: n = 7, 5 females; and WT_PD_: n = 7, 5 females). Mice were group-housed in ventilated cages under a 12/12h light/dark cycle with food and water available *ad libitum*. All experimental procedures were performed according to federal Swiss regulations under approval of the veterinary office of the Canton Zurich (license number ZH024/2021).

Mice were implanted as described before (Dias et al., n.d.; Kollarik et al., 2022; Morawska et al., 2021). Briefly, we placed 1 stainless steel screw per hemisphere, each located 2 mm posterior to Bregma and 2 mm lateral from midline. The screws were connected to a pin header via copper wires for electroencephalography (EEG) recordings, and the structure fixed with dental cement. For electromyography (EMG) recordings, we inserted two gold wires bilaterally into the neck muscles. All surgical procedures were performed under deep inhalation and local anaesthesia, with administration of anti-inflammatory and analgesic drugs. Postsurgical analgesia was administered for two days during both light and dark periods, with body weight and home cage activity daily monitored during the post surgical week and twice weekly thereafter (Dias et al., n.d.).

### EEG/EMG recordings and mCLAS paradigm

Following a two-week recovery, mice were individually housed in sound-insulated chambers, which continuously recorded EEG/EMG signals and delivered phase-locked auditory stimuli in real-time, as previously described (Dias et al., n.d.; Moreira et al., 2021). Briefly, we conducted inter-hemispheric tethered recordings (differential mode), with the left hemisphere EEG signal referenced to the right hemisphere electrode, amounting to one EEG channel per mouse (Dias et al., n.d.). All signals were sampled at 610.35 Hz, amplified after applying an anti-aliasing low-pass filter (45% of sampling frequency), synchronously digitized, and stored locally. We filtered real-time EEG between 0.1-36.0 Hz (2^nd^-order biquad filter), and EMG between 5.0-525.0 Hz (2^nd^-order biquad filter and 40 dB notch filter centered at 50 Hz). These signals informed the detection algorithms for stimuli delivery.

mCLAS was performed as described before (Dias et al., n.d.). Briefly, all mice received auditory stimuli consisting of clicks of pink 1/f noise (15ms, 35 dB SPL, 2ms rising/falling slopes) delivered by inbuilt speakers on top of the stimulation chambers. For phase-targeted mCLAS, a nonlinear classifier defined four decision nodes comprising power in EEG (EEGpower), power in EMG (EMGpower), a frequency component and a phase-target, which need to be simultaneously met to deliver sound stimuli, with a trigger delay of ∼29.30ms. The detailed computation of each node is described elsewhere (Dias et al., n.d.; Moreira et al., 2021). In WT and TG groups, we have previously defined 2Hz as the ideal frequency component to be tracked for both mouse strains used (Dias et al., n.d.). Regarding up-phase-targets, we set 30⁰ as optimal for the AD strain (WT_AD_ and AD mice) and 40⁰ for the PD strain (WT_PD_ and PD mice) (Dias et al., n.d.). We controlled for stimuli-evoked arousals or reflexes via in-built infrared recording cameras in each recording chamber. All triggers were flagged and saved for offline analysis. We recorded 2 EEG/EMG baseline days (no sound stimuli delivered but triggers flagged) and 2 EEG/EMG stimulation days for all mice.

### EEG/EMG offline scoring and pre-processing

We scored all recorded files using SPINDLE for animal sleep data (Miladinović et al., 2019), previously validated in our lab for mice (Dias et al., n.d.; Kollarik et al., 2022). SPINDLE retrieved 3 vigilance states (wakefulness: WAKE, non-rapid eye movement sleep: NREM sleep, and rapid eye movement sleep: REM sleep) with 4s epoch resolution from EEG/EMG recordings saved into EDF files (Browser™, TDT).

WAKE was characterized by high or phasic EMG activity for more than 50% of the epoch duration and low amplitude but high frequency EEG. NREM sleep was hallmarked by the presence of slow oscillations, reduced EMG activity and increased EEG power in the delta frequency band. REM sleep was defined based on high theta power and low EMG activity.

We pre-processed EEG data through a custom-written MATLAB script, as previously described (Dias et al., n.d.). In brief, we first removed artifacts by detecting clipping events, followed by a 3-point moving average and a basic Fermi window function, to prevent ringing artifacts (Dias et al., n.d.; Modarres et al., 2017). Next, we resampled the signal at 300Hz and filtered between 0.5–30Hz using low and high-pass zero-phased equiripple FIR filters. We inspected the signal for regional artefacts (2h sliding window) not detected during automatic scoring or the above mentioned steps (Dias et al., n.d.; Moreira et al., 2021). To pre-process EMG data, we first applied a basic Fermi window function, followed by resampling the signal at 300Hz and filtering between 5–100Hz using low and high-pass zero-phased equiripple FIR filters, with a bandstop centered at 50Hz (Dias et al., n.d.).

### EEG/EMG offline post-processing

#### <u>AD and PD strains</u>: Bout ratio and state space analysis (SSA)

We previously reported decreased NREM sleep fragmentation and microarousal levels upon mCLAS in AD mice (Dias et al., n.d.). To further assess vigilance state stability, we now calculated number of bouts per 24h, light period and dark period in WAKE, NREM sleep and REM sleep for each genotype of both the AD and PD strains, as a function of bout duration, categorizing bouts as short (0-40s) and long (>40s) relative to total bout duration in 24h (Kollarik et al., 2022). We then derived a ratio between the relative number of long and short bouts (bout ratio), at baseline (BL) and in stimulation (Stim). A higher bout ratio indicates a more consolidated vigilance state.

To gain further insight into the dynamics of vigilance states and their transitions, we performed a model-based EEG analysis comprising a two dimensional (2D) state space, based on previous literature on state space analysis (SSA) in rats, mice, healthy subjects and PD patients (Diniz Behn et al., 2010; Gervasoni et al., 2004; Imbach et al., 2012, 2016). The mathematical description of the model can be found elsewhere (Gervasoni et al., 2004; Imbach et al., 2012). In brief, we first performed spectral analysis of consecutive 4s EEG epochs (FFT routine, Hamming window, 2s overlap, 0.25Hz resolution) (Dias et al., n.d.). We then projected the spectral information of each epoch into a 2D plane by calculating two frequency ratios of predefined frequency bands adapted for mice and our pre-processing pipeline (ratio1: 6.5-9/0.3-9Hz; ratio2: 0.3-20/0.3-30Hz) (Diniz Behn et al., 2010). We further filtered the resulting time series with a 50-point moving average (Hann window) to avoid short-term fluctuations (Imbach et al., 2012). In this fashion, each epoch is represented as a single point in the 2D state space: a 24h recording is described as a scatterplot with clusters representing the different vigilance states (WAKE, NREM sleep and REM sleep), whereas trajectories in the 2D plane represent transitions between and within states. To quantify the dynamic changes across vigilance states upon mCLAS in the AD and PD strains, we calculated the Euclidean distances between vigilance state cluster centroids, during both light and dark periods (Imbach et al., 2016). Moreover, we calculated overall velocities within WAKE, NREM sleep and REM sleep in the state space, with velocity defined as the distance between two subsequent states divided by the time interval between the states (Diniz Behn et al., 2010; Imbach et al., 2012). We also calculated average state centroid distances and average overall velocities as percentage of change from BL values. Finally, we separated stable vigilance states (slow velocities) from transitions and fluctuations (fast velocities) by a heuristically defined velocity cut-off (v_limit_ = 0.02) based on visual inspection of the 24h distribution of all overall velocities (Imbach et al., 2012, 2016). This velocity-based approach allows to distinguish between slow velocities (v<0.02) which form consolidated clusters corresponding to vigilance states, from fast velocities (v>0.02) which correspond to transitions between vigilance states (Imbach et al., 2016). We calculated slow and fast velocities overall and per vigilance state (WAKE, NREM sleep and REM sleep) during the light and dark periods across the AD and PD strains, and quantified the number of slow and fast velocity events as percentage of change from BL values.

#### <u>AD strain</u>: Cumulative and hourly vigilance state proportions

To assess altered vigilance state patterns as a measure to probe a potential “sundowning” phenotype in the AD strain (Todd, 2020), we computed 24h, light period and dark period, cumulative and hourly vigilance state proportions as percentage of time spent in each state (WAKE, NREM sleep and REM sleep). Hourly proportions were calculated either 1) as an average of the hourly proportions of two baseline and two stimulation days across mice of the same genotype and strain (represented as light-to-dark proportions); or 2) as an average of the last baseline and the first stimulation day (represented as dark-to-light proportions) (Soltani et al., 2019). For each hourly proportion representation, we smoothed the data with a 3-point moving average (for visualization purposes) and computed the cumulative time spent per day in each vigilance state during the last 3h of the dark period. This time frame was chosen as it corresponds to the last hours of the active (dark) period in mice, immediately preceding the resting (light) period, mimicking the day-to-night (sundown) in humans (Bedrosian et al., 2011; Ognibene et al., 2005).

#### <u>PD strain</u>: EMG muscle tone, EEG slowing ratio, theta and alpha power

To assess possible loss of muscle atonia during REM sleep, as a measure to probe REM sleep behaviour disorder (RBD) in our PD strain (Schenck et al., 1986), we computed EMG mean muscle tone during each REM sleep bout, adapting an analysis pipeline previously described in rats (Garcia et al., 2017). Briefly, we first detrended the pre-processed EMG signal by removing the DC drift, rectified the resulting signal, and selected only REM sleep EMG bouts longer than 44s, excluding the first and last 4s of each bout to discard state transitions (Garcia et al., 2017). To allow for comparisons across days of recording and different animals, we normalized each REM sleep EMG bout to the maximum value of REM sleep in each day (REM sleep peak). Finally, we computed average normalized EMG muscle tone per 24h, light and dark periods, separately.

To attest for spectral power alterations, EEG general slowing and hyperexcitability in the PD strain (Cozac et al., 2016; Peters et al., 2020; Soikkeli et al., 1991), we extracted delta, theta, alpha and beta power through the spectral analysis described before for SSA, and normalized the data indicating the percentage of each bin with reference to the total spectral power between 0.5-30Hz, per 24h recording. Adapting the analysis pipeline of recent literature on EEG slowing in mild-cognitive impairment (MCI) patients (Lam et al., 2024), we computed an EEG slowing ratio defined as (δ+θ)/(α+β), with delta (0.5-4Hz), theta (4-10Hz), alpha (10-13Hz) and beta (13-20Hz), values derived from previous literature analysing EEG signal in this PD strain (Peters et al., 2020). Additionally, we calculated normalized average theta power in REM sleep across the different genotypes of the PD strain, with theta power values normalized to the maximum value of REM sleep in each day. Finally, we computed normalized average alpha power during WAKE and NREM sleep (power normalized to the maximum value of WAKE and NREM sleep in each day, respectively) as a proxy of cortical excitability, with lower power values corresponding to hyperexcitability (Pellegrino et al., 2024).

#### Statistical analysis

We conducted statistical analysis in IBM SPSS Statistics 28 (IBM, USA), with data plotted in GraphPad Prism 10.2.3 (GraphPad Software, LLC). We inspected the data for outliers through boxplots, with outliers considered as values 1.5 box-lengths from the edge of the box.

The distribution of each variable and its residuals was assessed for normality through skewness and kurtosis, while the homogeneity of variances was verified using Levene’s test with 5% significance. If normality and homogeneity of variances were met, one sample t-test, paired-sample t-test, two-sample t-test or one-way ANOVA with Šidák post hoc test were applied. Each test is reported with the corresponding p-value, unadjusted in case of single comparisons or adjusted in case of multiple comparisons, with significance set to 0.05. Results are shown as the mean and SEM.

## Results

### mCLAS enhances vigilance state stability in AD and PD mice

Alterations in vigilance state stability have been shown in neurodegeneration patients and animals (Kollarik et al., 2022; Lim et al., 2013). Whether a sleep enhancing approach based on slow-waves’ targeting is able to counteract such major structural deficiencies remains to be explored. We have preliminary shown decreased NREM sleep fragmentation and microarousal levels upon mCLAS in AD mice (Dias et al., n.d.). To assess vigilance state stability upon short-term mCLAS implementation and thus further probe mCLAS’ effect onto sleep maintenance and consolidation, we calculated the distribution of WAKE, NREM sleep and REM sleep bouts across genotypes of both the AD and PD strains, and calculated a long/short bout ratio for each vigilance state.

In AD mice, average baseline WAKE long/short bout ratio was significantly higher than in WT_AD_ controls (*p<0.05, two-sample t-test, **Figure 1A**) and lower than the ratio in AD mice after stimulation (**p<0.01, paired-sample t-test). Strikingly, baseline NREM sleep bout ratio in AD mice was significantly lower when compared to values of WT_AD_ controls (WT_AD_: 0.077±0.012; AD baseline: 0.038±0.07; *p<0.05, **Figure 1B**), with stimulation significantly increasing AD ratio towards control levels (**p<0.01). We observed no significant differences in REM sleep bout ratios across AD strain genotypes or treatments (**Figure 1C**).

**Figure 1.**
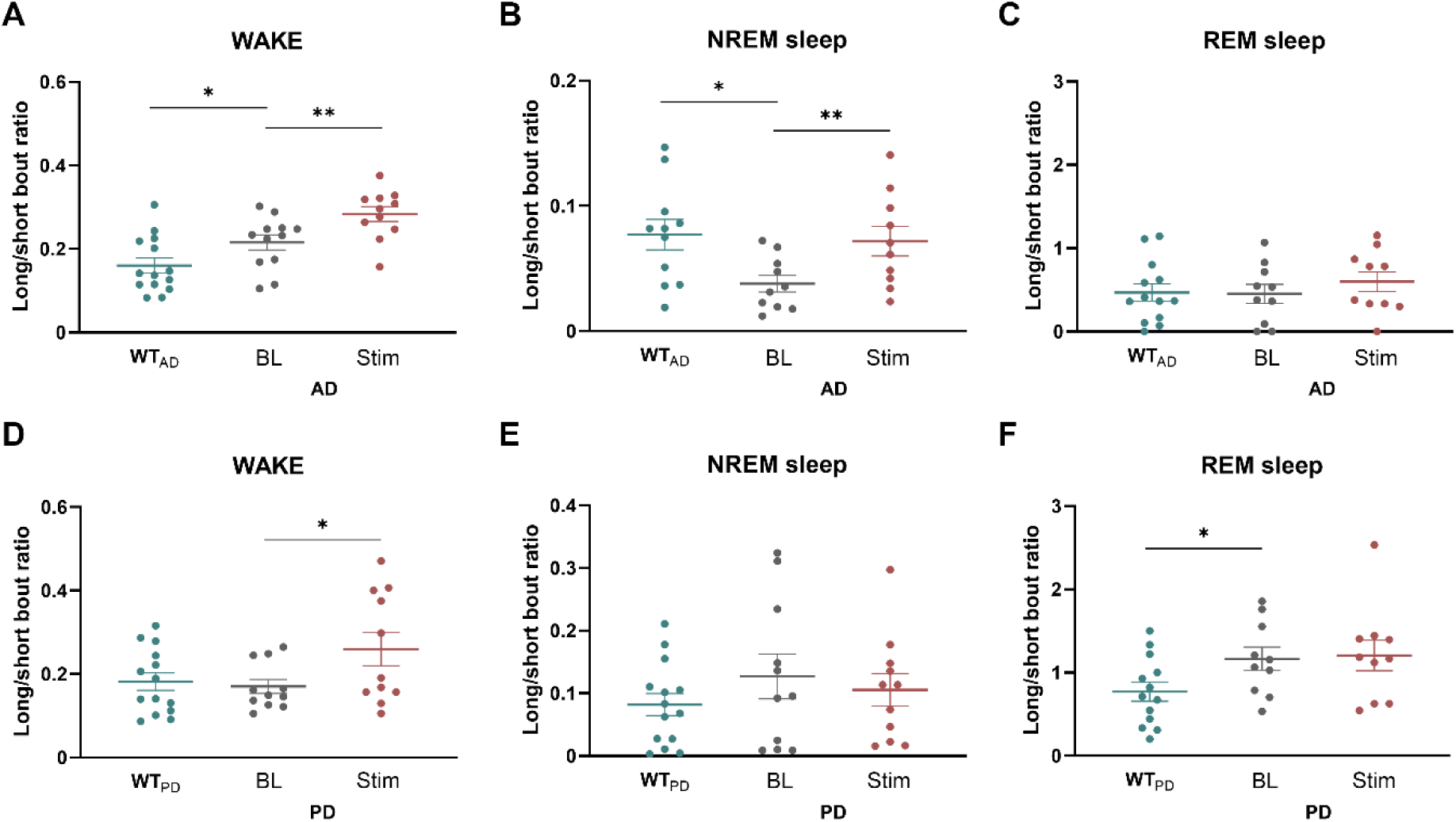
Long/short bout ratio per vigilance state, genotype and strain. **A:** Bout ratio during WAKE in WT_AD_, and in AD mice at baseline and after mCLAS. **B:** Bout ratio in the AD strain during NREM sleep. **C:** Bout ratio in the AD strain during REM sleep. **D:** Bout ratio during WAKE in WT_PD_, and in PD mice at baseline and after mCLAS. **E:** Bout ratio in the PD strain during NREM sleep. **F:** Bout ratio in the PD strain during REM sleep. Points in each scatter plot represent days from all mice, the full horizontal blue/grey/red lines the mean of the distributions and the whiskers the SEM. **p<0.01, *<0.05.

In PD mice, mCLAS significantly increased WAKE long/short bout ratio when compared to AD baseline values (*p<0.05, **Figure 1D**), which did not differ from WT_PD_ values. As counterpart, we observed a high NREM sleep long/short bout ratio in PD mice at baseline (0.128±0.036, **Figure 1E**) in relation to WT_PD_ ratio values (0.082±0.018), which decreased non-significantly towards WT_PD_ ratio values (0.10±0.02) upon mCLAS. Finally, PD baseline REM long/short bout ratio was significantly higher than WT_PD_ values (*p<0.05, **Figure 1F**), but was not significantly rescued upon mCLAS.

As both sleep and wake alterations and the specific effects of stimulation exerted onto them appeared distinct across the AD and PD models, we aimed for a more in-depth characterization of the changes in vigilance states’ dynamics and transitions across strains upon mCLAS. Accordingly, we performed state space analysis (SSA) (Diniz Behn et al., 2010; Imbach et al., 2012), projecting EEG spectral information of each vigilance state epoch onto 2D state space (**Figure 2A-2B**). Visual inspection of representative state space plots from WT_AD_ (**Figure 2A**) and AD mice (**Figure 2B**) hint distinctive vigilance state patterns between genotypes (note the scale difference between plots). We further characterized vigilance state proximity based on cluster centroid distances (coloured diamond shapes, **Figure 2A-2B**), and velocities of epoch trajectories within the 2D state space (**Figure 2C**).

**Figure 2.**
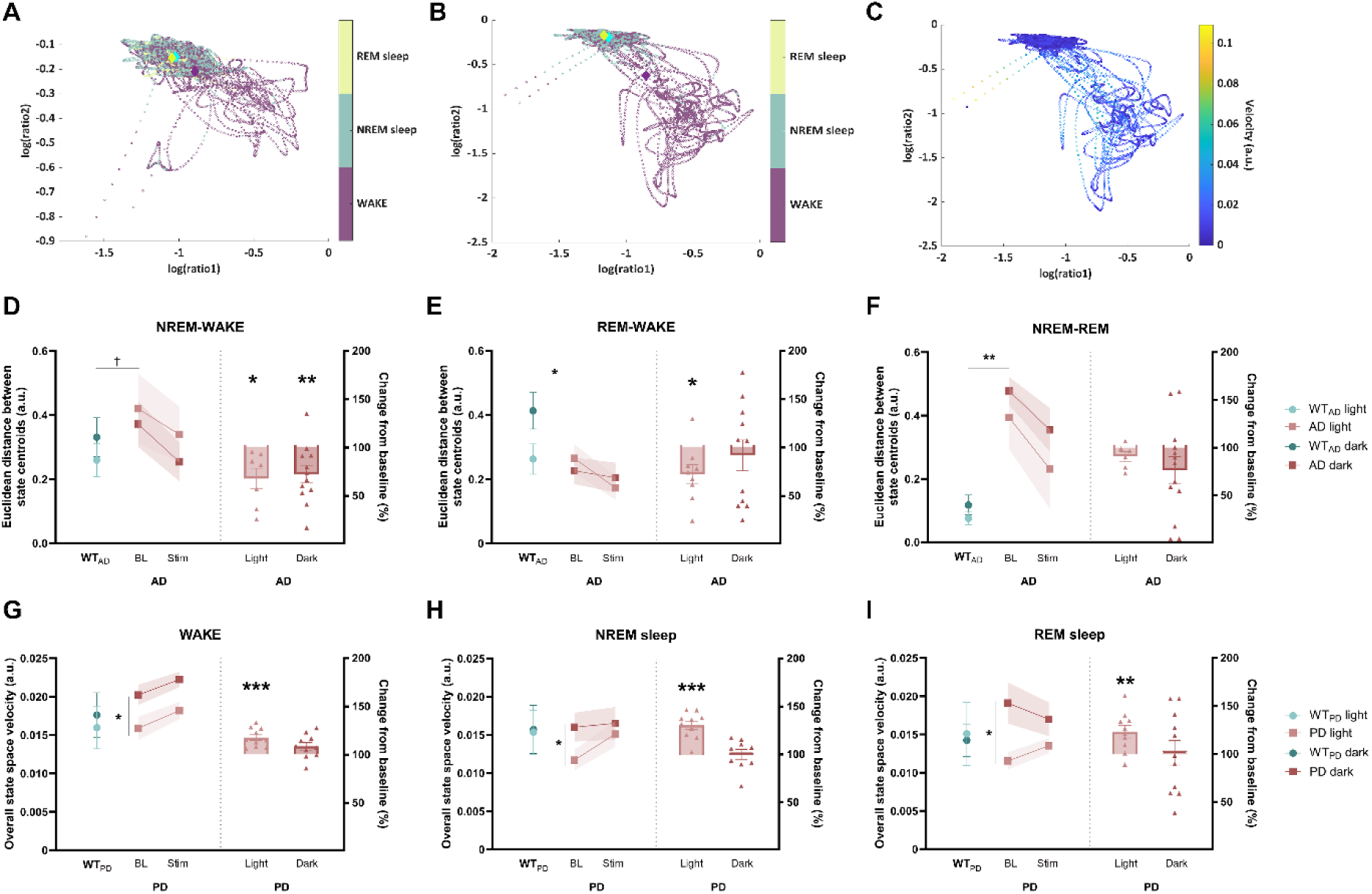
State space analysis (SSA), cluster centroid distances and trajectory velocities in the state space in AD and PD mice. **A:** Vigilance state patterns obtained via SSA analysis for a representative WT_AD_ mouse. **B:** Vigilance state patterns obtained via SSA analysis for a representative AD mouse (note the scale difference between 2A and 2B). Diamond coloured shapes represent cluster centroids for each vigilance state (Purple: WAKE; Blue: NREM sleep; Green: REM sleep). **C:** Velocities of epoch trajectories across the 2-D state space, for a representative AD mouse. Warmer/lighter colours represent fast velocities corresponding to vigilance state transitions, whereas colder/lighter colours represent slow velocities within vigilance states. **D-F:** Distances between NREM-WAKE, REM-WAKE and NREM-REM cluster centroids in AD mice, respectively. **G-I:** Overall state space velocities during WAKE, NREM and REM sleep in PD mice, respectively. Each before-and-after plot dot represents average 48h of baseline/stimulation Euclidean distance or velocity of each animal, and the shaded the SEM. In the bar plots, each scatter dot represents average 48h of Euclidean distance or state space velocity change as percentage of baseline of each animal, each bar the mean of such changes, and the whiskers the SEM. ***p<0.001, **p<0.01, *p<0.05.

Regarding vigilance state centroid distances, representing preserved or altered global sleep architecture (Imbach et al., 2016), average AD baseline NREM-WAKE distances appeared to be pathologically heightened in relation to WT_AD_ values, in both light and dark periods, decreasing to WT_AD_ level upon mCLAS (**Figure 2D, left**). Strikingly, NREM-WAKE centroid distances in AD mice decreased significantly as percentage of change from baseline values upon stimulation, in both light and dark periods (light: *p<0.05, one-sample t-test; dark: **p<0.01, **Figure 2D, right**). Moreover, REM-WAKE centroid distances in AD mice decreased significantly during the light period upon mCLAS (*p<0.05, **Figure 2E, right**) while no changes were elicited in the dark period. Interestingly, baseline NREM-REM centroid distances in AD mice were significantly higher than WT_AD_ values for both light and dark periods (**p<0.01, two-sample t-test, **Figure 2F, left**), decreasing non-significantly towards WT_AD_ levels upon mCLAS (**Figure 2F**). In PD mice, vigilance state centroid distances at baseline were not distinct from those of WT_PD_ mice nor changed significantly upon mCLAS (**Supplementary Figure 1A-1C**).

Regarding state space velocities, representing sleep state dynamics (Imbach et al., 2016), we found no significant differences in overall average velocities across vigilance states in AD mice at baseline compared to WT_AD_ mice nor upon stimulation (**Supplementary Figure 1D-1F**). Contrastingly, mCLAS’ effect onto overall state space velocities was striking in PD mice across all vigilance states (**Figure 2G-2I**). In WAKE, baseline overall velocities in PD mice in the light period appeared to be decreased in relation to PD baseline values in the dark period, increasing significantly upon mCLAS (***p<0.001, one-sample t-test, **Figure 2G**). Similarly, NREM sleep baseline overall velocities in PD mice during the light period were lower in relation to baseline dark PD values (*p<0.05, **Figure 2H, left**), increasing significantly upon mCLAS (***p<0.001, one-sample t-test, **Figure 2H, right**). We observed a similar effect in REM sleep, with baseline overall velocities decreased in PD mice during the light period and increased velocities during the dark period (*p<0.05, **Figure 2I, left**), with a significant increase of overall velocities during the light period towards WT_PD_ level upon mCLAS (**p<0.01, **Figure 2I, right**). Of note, WT_PD_ light and dark values are not significantly different throughout vigilance states (**Figure 2G-2I, left**), hinting that the observed baseline differences between light and dark velocities in PD animals are pathological.

Finally, we characterized separately state space slow velocities, corresponding to stable and consolidated vigilance states, from fast velocities, representing transitions and fluctuations between vigilance states (Imbach et al., 2016), within all vigilance states combined or WAKE, NREM sleep and REM sleep separately, during both light and dark periods across strains. To additionally have a measure of both consolidated and transitional events, we also quantified the number of slow and fast velocity events per vigilance state as percentage change from baseline values, respectively.

In AD mice, we observed no significant differences in overall slow velocities and number of slow velocity events either at baseline in comparison to WT_AD_ levels nor upon mCLAS for all vigilance states combined (**Supplementary Figure 2A-2B**). Analysis of slow velocities in each vigilance state separately, however, revealed a significant decrease in the abnormally high number of slow velocity events in AD mice towards WT_AD_ level upon mCLAS during REM sleep of both light and dark periods (light: **p<0.01; dark: *p<0.05, one-sample t-test, **Supplementary Figure 2C**). Analysis of fast velocities revealed that stimulation either significantly increased fast velocities during the light period in WAKE (*p<0.05, **Figure 3B**) or decreased fast velocities during NREM sleep (*p<0.05, **Figure 3C**), which did not differ from WT_AD_ levels at baseline. mCLAS’ effect on fast velocities in AD was particularly striking regarding the number of fast velocity events, which were significantly high at baseline during the dark period in comparison to light baseline AD values (*p<0.05, two-sample t-test, **Figure 3E, left**). Moreover, the number of fast velocity events during the light period significantly decreased across vigilance states in relation to baseline values (All vigilance states combined: ***p<0.001; WAKE: **p<0.01; NREM sleep: ***p<0.001; REM sleep: **p<0.001, **Figure 3E-3H, right**).

**Figure 3.**
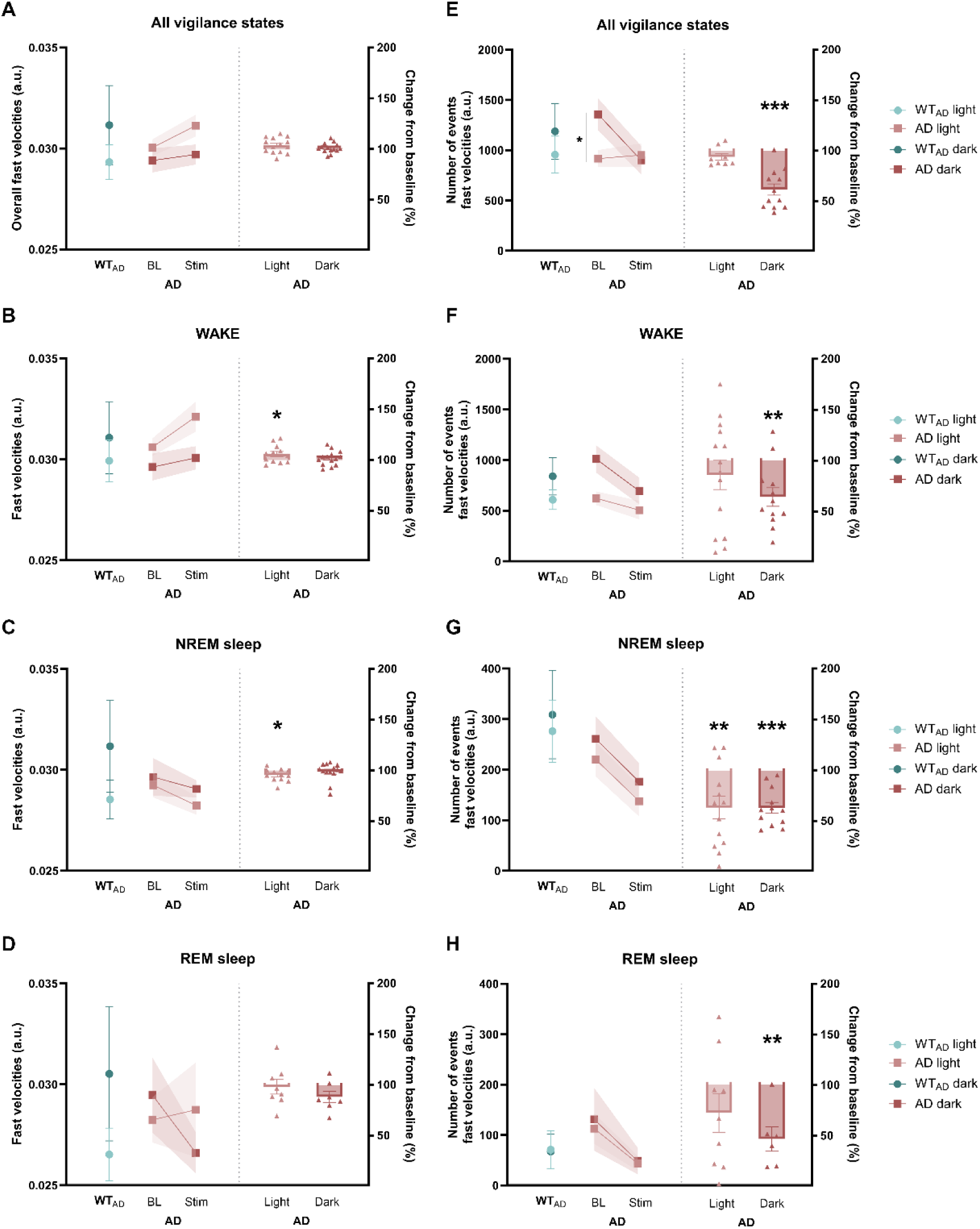
Fast state space velocities in AD mice. **A-D:** Fast velocities across all vigilance states, WAKE, NREM and REM sleep, respectively. **E-H**: Number of fast velocity events across all vigilance states, WAKE, NREM and REM sleep, respectively. Each before-and-after plot dot represents average 48h of baseline/stimulation Euclidean distance or velocity of each animal, and the shaded the SEM. In the bar plots, each scatter dot represents average 48h of Euclidean distance or state space velocity change as percentage of baseline of each animal, each bar the mean of such changes, and the whiskers the SEM. ***p<0.001, **p<0.01, *p<0.05.

In PD mice, we observed a significant decrease in the overall number of slow velocity events upon mCLAS during both light and dark periods (light: ***p<0.001; dark: **p<0.01, one-sample t-test, **Figure 4A**). In respect to slow velocities in each vigilance state upon mCLAS, we only observed a significant decrease in the abnormally high number of slow velocity events towards WT_PD_ levels during NREM sleep, within both light and dark periods (light: *p<0.05; dark: ***p<0.001, **Figure 4B**), and “faster” slow velocities during the dark period (*p<0.05). Regarding fast velocities, mCLAS significantly increased low fast velocities at baseline during both light and dark periods across all vigilance states (p-values between ***p<0.001 and *p<0.05, **Figure 4C-4F, left panels**). This striking effect in PD mice was accompanied by a significant increase in the number of fast velocity events during the light period across all vigilance states (All vigilance states: ***p<0.001; WAKE: ***p<0.001; NREM sleep: *p<0.05; REM sleep: *p<0.05, **Figure 4C-4F, right panels**).

**Figure 4.**
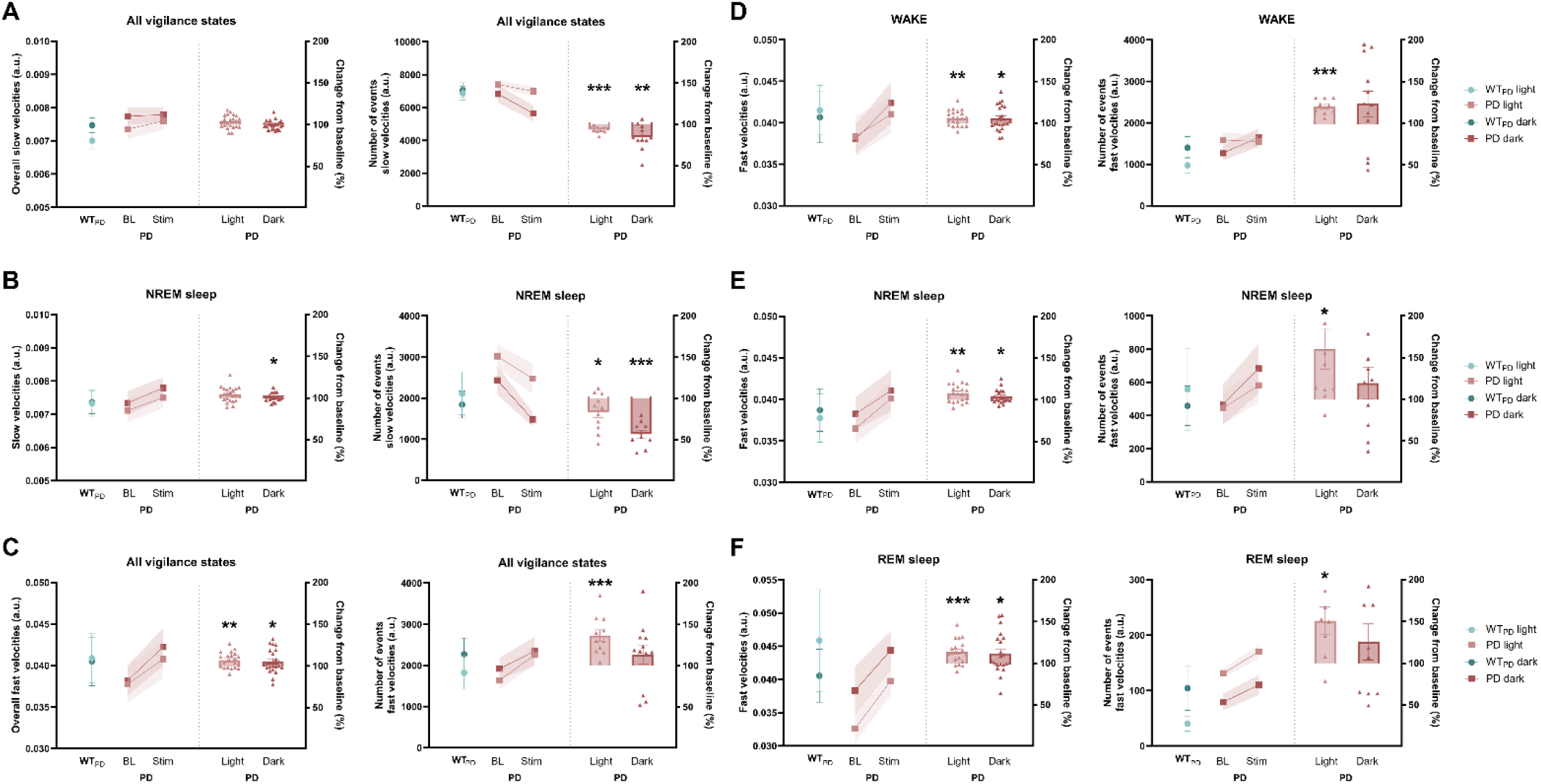
Fast state space velocities in PD mice. **A:** Slow velocities (left panel) and number of slow velocity events across all vigilance states (right panel). **B:** Slow velocities (left panel) and number of slow velocity events during NREM sleep (right panel). **C-F:** Fast velocities across all vigilance states, WAKE, NREM and REM sleep, respectively (**left panels**) and number of fast velocity events across all vigilance states, WAKE, NREM and REM sleep, respectively (**right panels**). Each before-and-after plot dot represents average 48h of baseline/stimulation Euclidean distance or velocity of each animal, and the shaded the SEM. In the bar plots, each scatter dot represents average 48h of Euclidean distance or state space velocity change as percentage of baseline of each animal, each bar the mean of such changes, and the whiskers the SEM. ***p<0.001, **p<0.01, *p<0.05.

Considering the dichotomous macrostructural sleep and wake phenotypes observed between the AD and PD models, as well as the selective modulation that mCLAS produces, we further separately explored mCLAS’ effect onto disease-specific sleep and wake alterations in each strain, with particular focus on hallmark sleep/wake impairments reported in patients with AD and PD.

### mCLAS counteracts sundowning-like phenotype in AD mice

Altered vigilance state patterns prior to sleep onset, increased locomotor activity, agitation and aggression, which are collectively described as “sundowning”, are common in patients with AD (Todd, 2020). To assess altered vigilance state patterns in the AD model, we calculated 24h, light and dark average proportions of WAKE, NREM sleep and REM sleep. To further explore vigilance states’ proportions dynamics, we analysed hourly percentages of time spent in each state (light-to-dark and dark-to-light representations, see Methods section), and derived the cumulative time spent during the last 3h of the active/dark period in each vigilance state (**Figure 5**).

**Figure 5.**
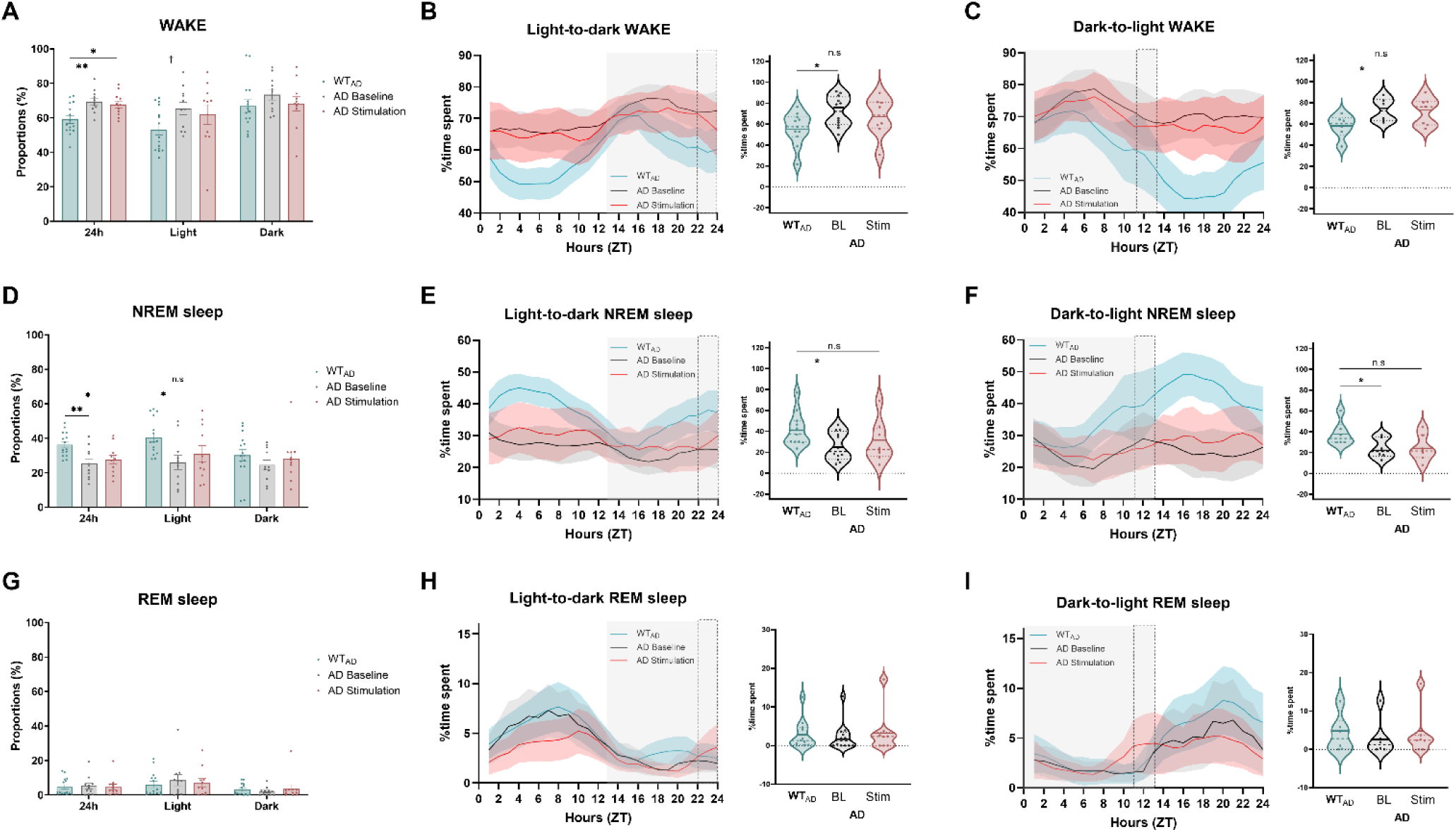
Vigilance state proportions in AD mice. **A, D and G**: WAKE, NREM and REM sleep proportions across 24h, light and dark periods, respectively. **B, E and H**: Light-to-dark representation of hourly vigilance state proportions, as percentage of time spent in each state (left panel), and cumulative time spent during the last 3h of the active/dark period (right panel) in WAKE, NREM and REM sleep, respectively. **C**: Dark-to light representation of hourly vigilance state proportions, as percentage of time spent in each state (left panel), and cumulative time spent during the last 3h of the active/dark period (right panel) in WAKE, NREM and REM sleep, respectively. In the bar plots, each scatter dot represents average daily proportions of each animal, each bar the mean, and the whiskers the SEM. In the 24h dynamics plots, blue/black/red shaded areas correspond to the SEM, the grey shaded panel corresponds to the dark period, and the dotted rectangle to the last 3h of the dark period. Points in each violin plot represent days from all mice, dashed and dotted lines the median and quartiles, and the full blue/black/red line the mean of the distributions. *p<0.05.

AD mice at baseline spent a significantly higher proportion of time in WAKE than WT_AD_ animals per 24h (**p<0.01, one-way ANOVA, Sidak correction, **Figure 5A**), which was not rescued by mCLAS (*p<0.05). Likewise, the cumulative time spent at baseline by AD mice in WAKE during the last 3h of the dark/active period was also higher than that of WT_AD_ animals, both in light-to-dark and dark-to-light representations (*p<0.05, **Figure 5B-5C**). In this specific time frame, the time spent in WAKE by AD mice undergoing mCLAS was non-significantly different from WT_AD_ values (**Figure 5B-5C**).

As a counterpart, AD mice at baseline spent a significantly lower proportion of time in NREM sleep than WT_AD_ animals per 24h (**p<0.01, one-way ANOVA, Sidak correction, **Figure 5D**), which was not rescued by mCLAS (*p<0.05). Contrastingly, even though at baseline AD mice spent a significantly lower proportion of time in NREM sleep than WT_AD_ animals during the rest/light period (*p<0.05), the time spent in NREM sleep was non-significantly different from WT_AD_ levels upon mCLAS (**Figure 5D**). Moreover, the cumulative time spent at baseline by AD mice in NREM sleep during the last 3h of the dark/active period was consistently lower than WT_AD_ animals, both in light-to-dark and dark-to-light representations (*p<0.05, **Figure 5E-5F**). In this time window, the time spent in NREM sleep by AD mice upon stimulation was non-significantly different from WT_AD_ values (**Figure 5B-5C**).We observed no significant differences between genotypes or upon treatment in REM sleep vigilance state proportions across 24h, light and dark periods, nor in the last 3h of the dark period (**Figure 5G-5I**).

### mCLAS counteracts cortical hyperexcitability phenotype in PD mice

REM sleep behaviour disorder (RBD) is considered a prodromal hallmark of PD, which is characterized by the loss of normal skeletal muscle atonia during REM sleep (Berg et al., 2021). Previous studies have shown that some PD rodent models recapitulate RBD-like phenotypes, with increased muscle tone during REM sleep (Garcia et al., 2017; Okuda et al., 2022; Taguchi et al., 2020). To assess alterations in muscle tone in the PD model, we separately computed average normalized EMG muscle tone during REM sleep per 24h, light and dark periods (**Figure 6**). In contrast to previous data, PD mice at baseline displayed significantly lower muscle tone than WT_PD_ mice during REM sleep across 24h, light and dark periods (24h: *p<0.05; light:*p<0.05; dark: † p=0.06, two-sample t-test, **Figure 6A-6C**), suggesting the presence of abnormal REM sleep muscle hypotonia in this model. Upon mCLAS, muscle tone in PD mice across 24h, light and dark periods remained significantly different from WT_PD_ levels (24h: *p<0.05; light:**p<0.01; dark: *p<0.05).

**Figure 6.**
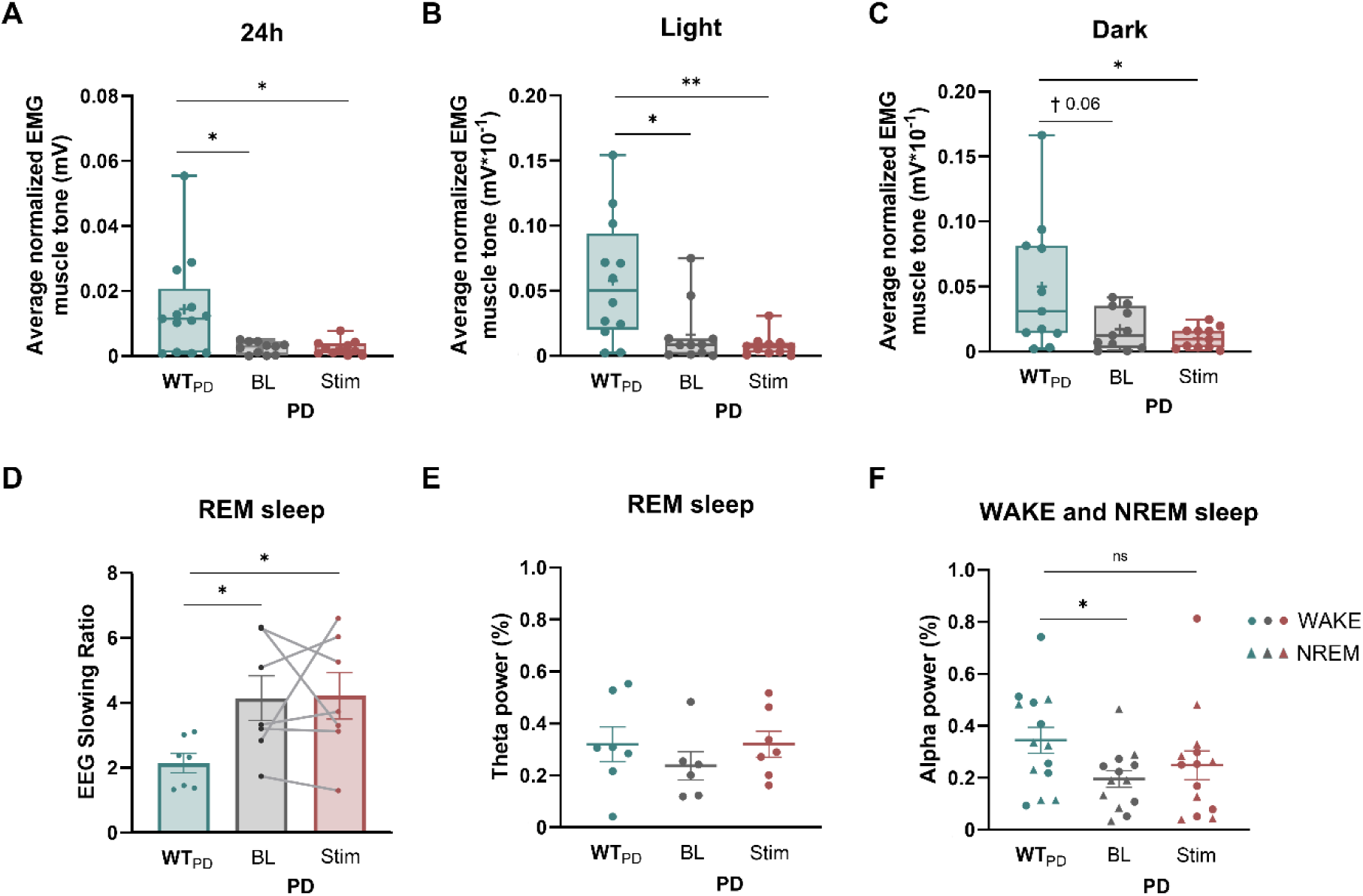
EMG muscle tone, EEG slowing ratio, theta and alpha power in PD mice. **A-C:** Average normalized EEG muscle tone during REM sleep in 24h, light and dark periods, separately. **D:** EEG slowing ratio calculated during REM sleep. **E:** Average normalized theta power during REM sleep. **F:** Average normalized alpha power during WAKE and NREM sleep. In the box-plots, the dots represent average daily muscle tone values from all mice, the edges of the box are the 25^th^ and 75^th^ percentiles, the whiskers extend to the minimum and the maximum of the distribution, the full line the median and the ‘+’ the mean. In the bar plots, each scatter dot represents average slowing ratio of each animal, the connecting lines represent before-and-after for the same mouse, each bar the mean, and the whiskers the SEM. Points in each scatter plot represent power in WAKE (circle) and NREM sleep (triangle) for each mouse, the full horizontal blue/grey/red lines the mean of the distributions and the whiskers the SEM. **p<0.01, *p<0.05, †p<0.1.

Generalized slowing in the EEG of PD patients has also been reported as an hallmark of PD (Soikkeli et al., 1991), and has been identified in M83 mice (Peters et al., 2020), with altered power values in delta, theta, alpha and beta frequencies. To attest for generalized EEG slowing, we computed an EEG slowing ratio adapted from human sleep literature (Lam et al., 2024). As expected, we observed a significantly increased EEG slowing ratio in PD mice at baseline compared to WT_PD_ mice (*p<0.05, two-sample t-test, **Figure 6D**), alteration that remained present upon mCLAS (*p<0.05).

Spectral theta power is also impaired in prodromal PD patients (Cozac et al., 2016; Zhu et al., 2019). To explore spectral theta power alterations in the PD model at baseline and upon mCLAS, we calculated average theta power during REM sleep. Theta power values were non-significantly different in PD mice at baseline in relation to WT_PD_ values, which increased non-significantly upon mCLAS (**Figure 6E**). Additionally, M83 transgenic mice have also been previously reported to display hyperexcitability during WAKE and NREM sleep (Peters et al., 2020), linked to decreased alpha power (Pellegrino et al., 2024). We thus calculated average alpha power during WAKE and NREM sleep combined, finding that alpha power values in PD mice at baseline were significantly lower in relation to WT_PD_ values (WT_PD_: 0.34±0.05%; PD baseline: 0.19±0.03%, *p<0.05, two-sample t-test, **Figure 6F**). Alpha power values in PD mice upon mCLAS were indistinct to those of WT_PD_ mice.

## Discussion

In this study, we show that up-phase-targeted mCLAS rescues some key pathological sleep traits in two mouse models of neurodegeneration. To achieve this, we used mCLAS parameters previously optimised for each disease model in a 2-day stimulation design, and conducted an in-depth analysis of the acute effects of mCLAS onto sleep macro- and microstructure, and disease-specific sleep and wake disturbances previously described in patients and animal models. Strikingly, mCLAS selectively prompted the acute alleviation of some neurodegeneration-associated sleep and wake phenotypes, by either potentiating sleep maintenance and stability in AD mice or by rescuing bradysomnia and cortical hyperexcitability in PD animals.

### mCLAS rescues altered sleep maintenance and stability in AD mice

Decreased sleep continuity is a distinctive feature of early-stage AD (Bubu et al., 2017; Lim et al., 2013; Sindi et al., 2018), being linked to cognitive impairment in patients, accelerated microglia and astrocyte aging and activation, increased amyloid-beta deposition, tau phosphorylation, neuroinflammation, and decreasing AQP4 expression in AD animals (Cai et al., 2019; Kaneshwaran et al., 2019; Vasciaveo et al., 2023). Moreover, prodromal AD mice of our strain were reported to exhibit heightened NREM sleep fragmentation accompanied by a decrease in NREM sleep duration compared to healthy controls (Kollarik et al., 2022; Wisor et al., 2005).

Our short-term up-phase targeted mCLAS paradigm appears to restore a pathologically low ratio between long and short NREM sleep bouts in AD mice towards healthy WT_AD_ levels, perhaps indicating increased sleep stability. These results corroborate previous data showing a 14% decrease in NREM sleep fragmentation index and decreased microarousal events upon mCLAS in AD mice (Dias et al., n.d.). Moreover, a striking effect of the mCLAS paradigm was prompting the decrease of pathologically heightened distances between vigilance states within the 2D state space, again suggesting a more consolidated sleep architecture in these animals (Imbach et al., 2012, 2016). In addition, mCLAS decreased the high number of transitions in all vigilance states, hinting on improved sleep microstructure (Diniz Behn et al., 2010; Imbach et al., 2012, 2016). The current results confirm our previous speculation that mCLAS corrected these sleep impairments in AD mice via impeded NREM-REM and NREM-WAKE transitions, potentiating sleep consolidation and maintenance (Dias et al., n.d.; dos Santos Lima et al., 2019; Gradwohl et al., 2017). Faster sleep dissipation as a consequence of higher neuronal population synchrony is also a phenomena that mCLAS might have enhanced (Bernardi et al., 2018; Fattinger et al., 2014; Krugliakova et al., 2022; Vyazovskiy et al., 2009), which shall be verified in future studies.

Other hallmarks of prodromal AD include altered body temperature and hormone release, increased locomotor activity and behavioural disruptions in the late evening (before bedtime), described as the sundowning circadian syndrome in AD patients (Todd, 2020). Altered vigilance state patterns, locomotor activity and body temperature related with sundowning disturbances were also described in mouse models of AD, particularly in the transition between the dark (active) and light (rest) periods (Bedrosian et al., 2011; Ognibene et al., 2005; Warfield et al., 2023).

Verifying previous reports on altered sleep proportions in AD mice (Kollarik et al., 2022), our mutant animals displayed a significantly higher amount of time spent in WAKE and lower amount of time spent in NREM sleep at baseline when compared to WT_AD_ mice. We observed the same pathological patterns when focusing solely on the last 3h of the dark period (the active phase in mice), corresponding to late afternoon sundowning symptoms’ window of humans, which aligns with previous reports on increased locomotor activity during this period in this AD model (Bedrosian et al., 2011; Ognibene et al., 2005; Oyegbami et al., 2017). Moreover, hourly 24h dynamics of the percentage of time spent in each vigilance state show flattened activity rhythms in AD mice at baseline, which are consistent with previous reports in patients and animals models of AD (Bedrosian et al., 2011; Harper et al., 2005). Notably, mCLAS seems to have counteracted the sundowning phenotype observed in WAKE and NREM sleep in the last 3h of the dark period, with 24h hourly dynamics acquiring a less flattened pattern towards WT_AD_ levels. These results indicate that short-term mCLAS may have an effect onto circadian sleep proportions in AD mice, which is sufficient to lessen vigilance states’ disturbances without fully restoring physiological healthy levels. It is conceivable that the acute nature of our protocol could be responsible for this partial recovery, suggesting that a cumulative mCLAS effect over a long-term paradigm might be necessary to fully reset AD-triggered sleep and wake disturbances.

### mCLAS rescues bradysomnia and cortical hyperexcitability in PD mice

Similarly to other neurodegenerative diseases, sleep disturbances in PD patients include insomnia, increased sleep fragmentation and nocturnal arousals, excessive daytime sleepiness (EDS), slowing of patients’ EEG signal and abnormalities in spectral EEG frequencies (Bollu & Sahota, 2017; Cozac et al., 2016; Dodet et al., 2024; Lam et al., 2024; Menza et al., 2010; Soikkeli et al., 1991; Zhu et al., 2019).

In line with our previous observation of a lack of abnormal sleep fragmentation in the PD model, long/short WAKE and NREM sleep bout ratios were not impaired at baseline in PD mice when compared to WT_PD_ animals (Dias et al., n.d.; Morawska et al., 2021). However, our mCLAS paradigm significantly increased the ratio between long and short WAKE bouts, which were not significantly altered at baseline when compared to WT_PD_ mice. Additionally, mCLAS seems to decrease NREM sleep bouts’ length, which were non-significantly heightened in PD mice at baseline in relation to WT_PD_ mice. These initially puzzling results become apparent through our more in-depth state space analysis: mCLAS acts on over consolidated NREM sleep states in PD mice by decreasing the number and speed of slow velocities (within state), which were abnormally high at baseline. Strikingly, mCLAS prompts an increase in transition speed and occurrence across all vigilance states, with transition speed being impaired at baseline in PD mice, as previously reported in PD patients and termed *bradysomnia* (Imbach et al., 2016). As formerly hypothesized, lower transition velocity is already observed at a prodromal PD stage in our mice, raising the urge to consider bradysomnia as a potential early disease biomarker (Imbach et al., 2016). Prompting of faster state transitions towards healthy levels as well as benefits in subjective sleep quality were previously observed in PD patients upon dopaminergic treatment (Brunner et al., 2002; Imbach et al., 2016; Trenkwalder et al., 2011). Whether mCLAS rescued altered sleep-wake dynamics in PD mice by counteracting dopaminergic neurotransmission deficits remains to be directly explored. Moreover, M83 mice have been previously reported to lack striatal tyrosine hydroxylase (enzyme catalyzing L-DOPA, a precursor of dopamine) at a prodromal age, as well as displaying increased D1 receptors in the substantia nigra and decreased dopamine transporters in the nucleus accumbens and striatum (Pupyshev et al., 2018; Unger et al., 2006), suggesting that dopamine neurotransmission deficiencies exist in this model and may have been targeted by neurotransmission processes under the broad influence of CLAS-enhanced deep sleep. Testing this hypothesis is not within the scope of this study but should be further explored via experimental designs coupling mCLAS with quantitative methods such as in vivo microdialysis, liquid chromatography or imaging techniques (Jensen et al., 2017; Post & Sulzer, 2021).

Additionally, our mCLAS paradigm was also able to rescue baseline pathologically low alpha power during WAKE and NREM sleep in PD mice. Alpha power is widely considered a marker of the Excitation/Inhibition balance (E/I), with low alpha power corresponding to high E/I (Pellegrino et al., 2024). Consequently, low alpha power is indicative of cortical hyperexcitability, which is observed in both M83 mice and in prodromal PD patients (Pellegrino et al., 2024; Su et al., 2023). Thus, mCLAS seems to modulate intrinsic cortical excitability, perhaps upon manipulation of the E/I balance by potentiating either decreased excitatory (glutamatergic) or increased inhibitory (GABAergic) synaptic transmission (Kirischuk, 2022; Pellegrino et al., 2024).

Slowing of EEG signal, conceived to represent the loss of cells along ongoing neurodegeneration, is commonly observed in PD patients (Soikkeli et al., 1991) for which specific measures have been developed in humans to quantify it (Lam et al., 2024). Adapting those measures to our experimental conditions, we did not observe mCLAS-triggered changes in the enhanced EEG slowing ratio observed in PD mice, confirming the not surprising scenario that this short-term (2-day) mCLAS intervention was not able to counteract the neurodegenerative process. Whether long-term mCLAS could impact disease pathophysiology thus preventing cell loss and consequently counteract EEG slowing will be explored in future studies. Nevertheless, we do not exclude the possibility that this human-derived EEG slowing metric is perhaps not ideal to probe general slowing is our experimental conditions: even though mCLAS increases alpha power, this paradigm also elicits an increase in theta (non-significantly) and delta power, the later verified in our previous report (Dias et al., n.d.). Further studies should explore new EEG slowing assessment metrics accounting for the nature of mCLAS interventions.

Finally, the loss of skeletal muscle atonia during REM sleep in RBD is also described as an important underpinning of prodromal PD (Berg et al., 2021; Postuma, 2014; Schenck et al., 1986). Therefore, based on previous literature exploring RBD in rats (Garcia et al., 2017), we determined EMG muscle tone during REM sleep in the PD model to explore whether mCLAS might have influenced EMG activity as shown before via modulated microarousal activity in AD mice (Dias et al., n.d.). Surprisingly, the muscle tone of PD mice at baseline was significantly decreased when compared to WT_PD_ levels, with mCLAS exerting no significant changes. Increased atonia during REM sleep in PD mice at baseline might act as a compensation mechanism for the increased epileptic seizures observed during WAKE and NREM sleep in our strain (linked with cortical hyperexcitability), but this speculation should be further probed (Peters et al., 2020). To the best of our knowledge, there are currently no accounts of RBD-like symptoms in our PD strain, hinting that this model might not fully recapitulate RBD symptoms in PD patients, with the need to recur to other PD models to observe this sleep phenotype (Okuda et al., 2022; Taguchi et al., 2020).

### mCLAS selectively prompts an adaptive rescue of disease-specific sleep and wake alterations across strains

Our mCLAS paradigm appears to adaptively prompt a repair towards healthy levels, by either potentiating sleep consolidation and vigilance state stability in AD mice or rescuing bradysomnia and decreasing cortical hyperexcitability in PD mice. We have previously observed a similar specific effect regarding mCLAS’ modulation of NREM sleep fragmentation and microarousal events, both altered in AD but not in our PD strains at baseline (Dias et al., n.d.). Moreover, we have also reported on strain specificity regarding parameter optimization for SWA enhancement via phase-locked mCLAS, suggesting that genetic background might have a significant role when tailoring this neuromodulation technique to clinical populations. The present results show that baseline sleep impairments differed across neurodegeneration strains, determining mCLAS’ effects in counteracting the disease-specific sleep disturbances observed. Hence, we conclude that mCLAS allows for a highly specific alleviation of neurodegeneration-associated sleep and wake phenotypes, correcting only existing baseline impairments adaptively across strains. These combined results are highly relevant for CLAS in human studies and future clinical implications, suggesting that when tailored to a certain neurodegenerative disease, CLAS could indeed be a promising candidate as a preventive and/or therapeutic tool to specifically rescue vigilance states’ alterations bound to that disease. Our encouraging results provide evidence for acute disease-specific sleep and wake benefits prompted by mCLAS in two neurodegeneration models, with long-term studies still needed to further evaluate potential rescue of behavioural and neuropathological alterations upon stimulation.

### Limitations

We have not performed sex-based analysis in this study, with the results including a higher percentage of female individuals due to male dropout based on aggressiveness. Data loss in several mice also prevented repeated measures analysis at times. This caveat was mainly due to software crashes, lost EEG implants, disconnections or poor signal quality associated with prolonged recordings. In terms of experimental design, our short-term intervention did not allow to assess the long-term effects of mCLAS onto sleep disturbances, cognition or pathological protein burden, for which longer stimulation protocols might be needed (Zeller et al., 2024). Lastly, we did not implement adaptive feedback methods to estimate ongoing phase-targets in our experiments (Hebron et al., 2024; Navarrete et al., 2020; Papalambros et al., 2019), which could potentially improve applicability of long-term mCLAS during deep sleep.

## Conclusions

In summary, we show that key pathological sleep-wake traits are adaptively rescued by mCLAS in AD and PD mice models, prompting the acute alleviation of neurodegeneration-associated sleep phenotypes. The present results pave the way for experiments unravelling the mechanisms underlying mCLAS’ modulation of SWA, its potential long-term effect on brain disease, and the role of SWA in toxic protein accumulation processes in neurodegeneration. Hence, mCLAS may become the future cornerstone of SWA-based therapies for neurodegenerative diseases.

## Author contributions

**Inês Dias**: Conceptualization, Methodology, Software, Validation, Formal Analysis, Investigation, Data Curation, Visualization, Funding acquisition, Writing – Original Draft, Writing – Review & Editing; **Christian R. Bauman**: Funding acquisition, Writing – Review & Editing; **Daniela Noain**: Conceptualization, Resources, Supervision, Project administration, Funding acquisition, Writing – Original Draft, Writing – Review & Editing.

## Acknowledgments and funding statement

We would like to thank Ami Beuret and Lukas Imbach for their valuable support. This project was funded by Parkinson Schweiz (ID, DN), the Dementia Research Switzerland - Synapsis Foundation via an earmarked donation of the Armin and Jeannine Kurz Stiftung (DN), the Dr. Wilhelm Hurka Foundation (DN), and the Swiss National Science Foundation (DN). The funding sources had no involvement in study design, collection, analysis, interpretation of the data nor in writing or deciding to submit this article for publication.

## Supplementary figures

**Supplementary Figure 1.**
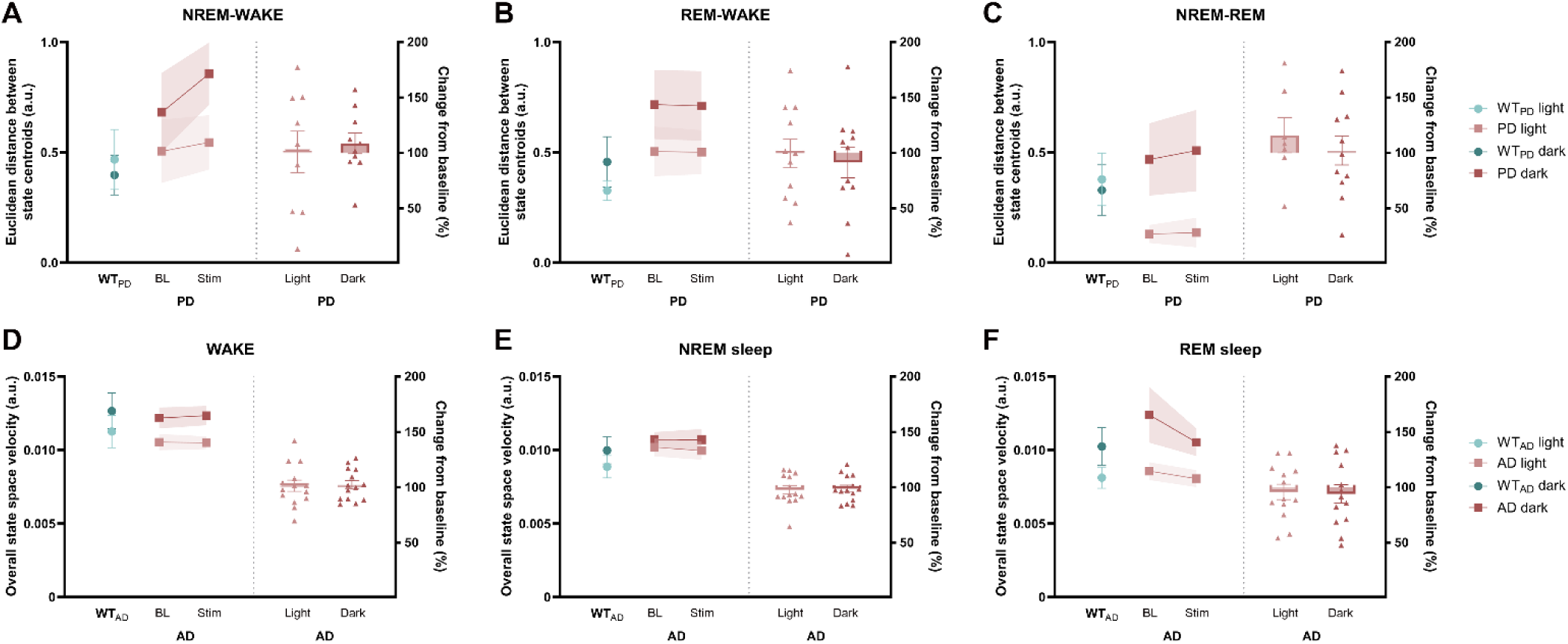
Cluster centroid distances in PD mice and trajectory velocities in the state space in AD mice. **A-C:** Distances between NREM-WAKE, REM-WAKE and NREM-REM cluster centroids in PD mice, respectively. **D-F:** Overall state space velocities during WAKE, NREM and REM sleep in AD mice, respectively. Each before-and-after plot dot represents average 48h of baseline/stimulation Euclidean distance or velocity of each animal, and the shaded the SEM. In the bar plots, each scatter dot represents average 48h of Euclidean distance or state space velocity change as percentage of baseline of each animal, each bar the mean of such changes, and the whiskers the SEM.

**Supplementary Figure 2.**
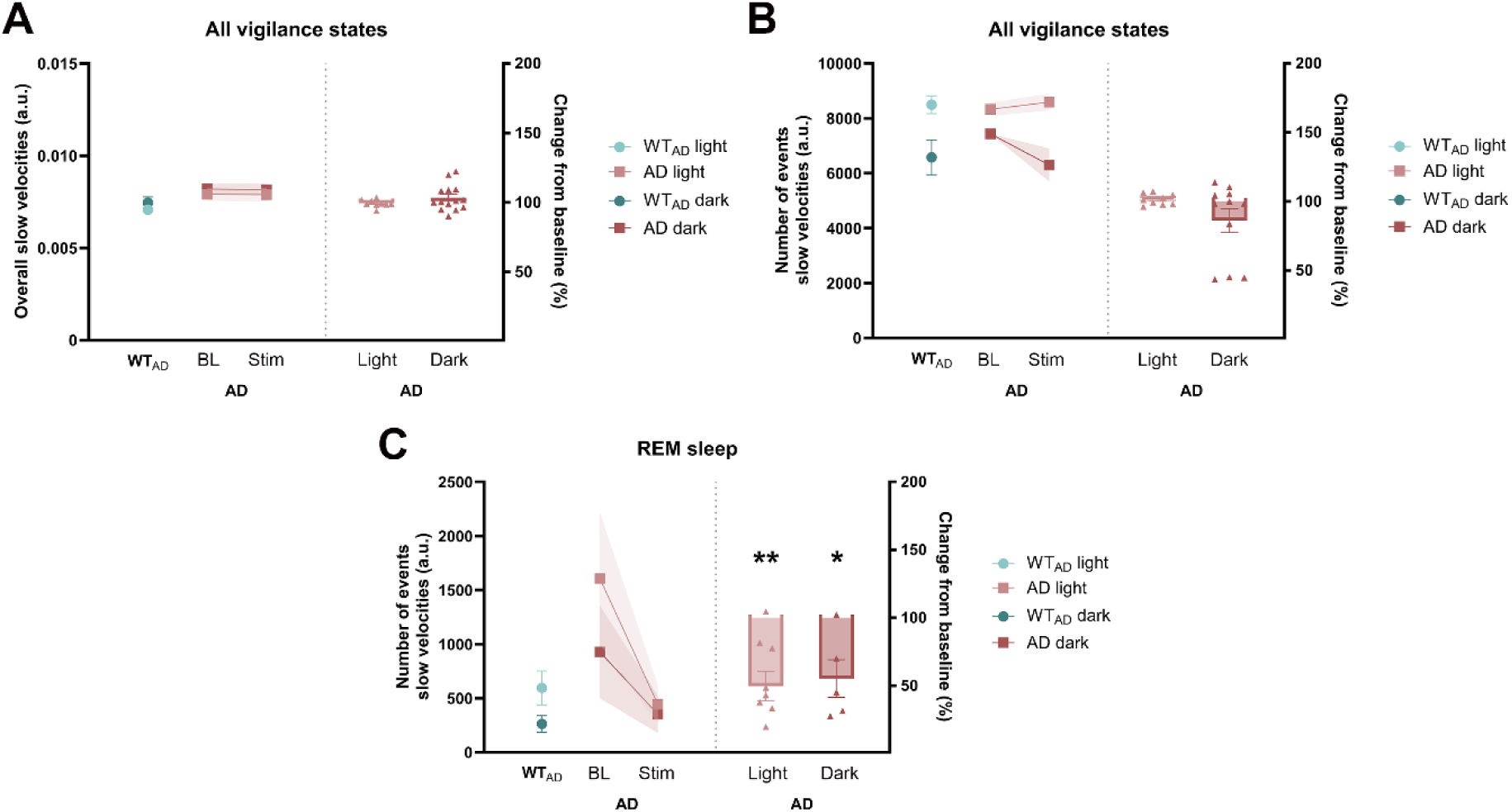
Slow state space velocities in AD mice. **A:** Slow velocities across all vigilance states. **B:** Number of slow velocity events across all vigilance states. **C:** Number of slow velocity events during REM sleep. Each before-and-after plot dot represents average 48h of baseline/stimulation Euclidean distance or velocity of each animal, and the shaded the SEM. In the bar plots, each scatter dot represents average 48h of Euclidean distance or state space velocity change as percentage of baseline of each animal, each bar the mean of such changes, and the whiskers the SEM. **p<0.01, *p<0.05.

